# Concurrent TMS and Spinal Cord fMRI Reveal Intensity-Dependent Modulation of Spinal Circuits

**DOI:** 10.64898/2026.07.24.740278

**Authors:** Ekansh Sareen, Rebecca Jones, Estelle Raffin, Friedhelm C. Hummel, Dimitri Van De Ville

## Abstract

The spinal cord serves as a crucial relay for motor commands, yet the role of its local circuitry in sensorimotor integration remains poorly understood. Most non-invasive cortical stimulation studies, rely on electrophysiological readouts or inferred spinal function from corticospinal anatomy, leaving the downstream impact of cortical stimulation on spinal circuitry largely uncharted in vivo. Advances in spinal cord functional MRI (SC-fMRI) now enable spatially resolved imaging of segmental gray and white matter and their interactions with descending cortical inputs. Here, we introduce a multimodal framework that combines single-pulse transcranial magnetic stimulation (TMS) of the primary motor cortex with SC-fMRI to probe TMS-evoked spinal activity in humans. Using graded TMS intensities, we examined blood oxygenation level-dependent (BOLD) responses in the cervical spinal cord and asked how spinal activation depends on effective engagement of the descending motor system. Our findings reveal robust, intensity-dependent spinal BOLD responses aligned with descending pathways, with activation concentrated in expected territories such as the lateral corticospinal tract and ventral horn at segments innervating the stimulated hand muscle. By linking peripheral output to segment- and pathway-resolved spinal signals, these results demonstrate that concurrent TMS–SCfMRI can capture, in vivo, how cortical drive is expressed within human spinal circuitry and provide a new framework to measure spinal contributions to sensorimotor control with spatial specificity beyond traditional peripheral readouts.

**Graphical Abstract:** 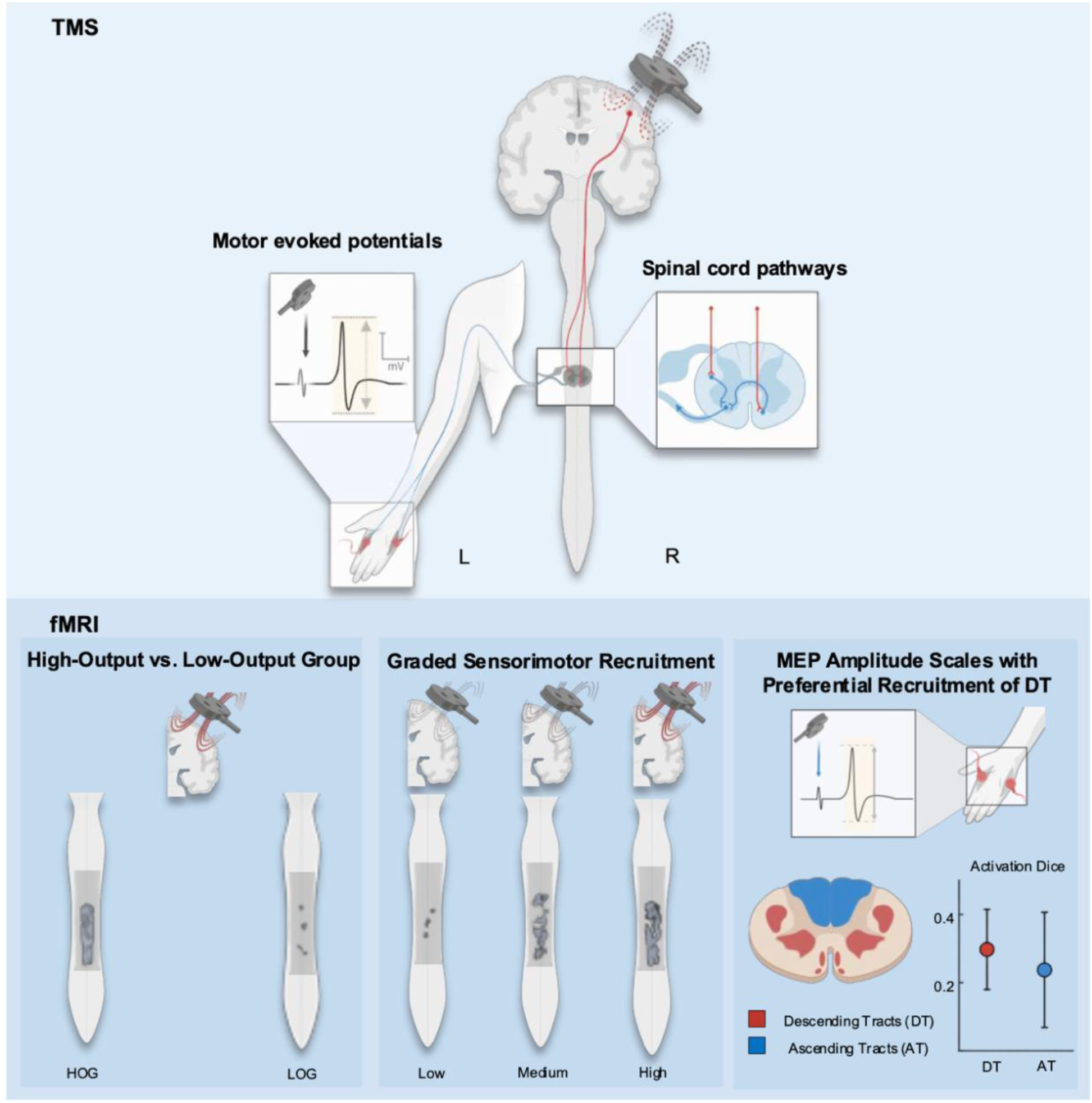

## Introduction

Functional magnetic resonance imaging (fMRI) has been central to advancing our understanding of the central nervous system (CNS). By leveraging blood oxygenation level-dependent (BOLD) contrast, fMRI provides a non-invasive measure of neuronal activity through associated hemodynamic responses (Logothetis et al., 2001) and has been widely applied in task-based and resting-state paradigms to map functional brain networks (van den Heuvel & Pol, 2010). In contrast to the substantial body of work focusing on the brain, the spinal cord, despite being an integral part of the CNS and a key mediator of sensorimotor functions (Darby & Frysztak, 2014), has remained comparatively underexplored using fMRI. This discrepancy reflects the anatomical and physiological challenges of spinal imaging, in which the small cross-sectional area, proximity to osseous structures, magnetic field inhomogeneities, and physiological noise from cardiac and respiratory cycles degrade signal quality (Giove et al., 2004; Stroman et al., 2014). Despite these limitations, recent innovations in acquisition strategies (Combes et al., 2022; Landelle et al., 2021) and post-processing methodologies (Eippert et al., 2017; De Leener et al., 2017; Kinany et al., 2023), including signal processing approaches such as innovation-driven co-activation patterns (iCAPs) based on the Total Activation framework (Karahanoğlu & Van De Ville, 2015; Karahanoğlu et al., 2013), have made spinal fMRI increasingly feasible and reliable (Wheeler-Kingshott et al., 2014). In particular, methods such as iCAPs apply regularized hemodynamic deconvolution to estimate sustained activity-inducing signals and have been used to identify spatially coherent, functionally meaningful resting-state patterns in the spinal cord (Kinany et al., 2020). Their efficacy to enhance sensitivity to transient and subtle activation changes has been systematically validated in spinal cord fMRI (Sareen et al., 2025), supporting their use to improve detection power in task-based paradigms. Building on these advances, task-based spinal fMRI paradigms have revealed region-specific activations that align with established neuroanatomical principles, with sensory stimulation engaging dorsal horn regions and motor tasks recruiting ventral horn territories (Landelle et al., 2021), while more complex patterns, including dorsal-ventral overlap and co-activation during both sensory and motor tasks (Landelle et al., 2021), point to integrative sensorimotor processing that likely reflects descending cortical inputs and local inter-neuronal circuits (Willis & Coggeshall, 2012). This interplay between top-down and local spinal mechanisms highlights the need for approaches that can selectively modulate and probe distinct components of spinal circuitry.

In this context, noninvasive neurostimulation techniques such as transcranial magnetic stimulation (TMS) offer targeted modulation of neuronal circuitry. TMS delivers brief, high-intensity magnetic pulses that generate electric fields penetrating the scalp and skull, inducing localized currents in cortical neurons and modulating neural excitability in a controlled and repeatable manner (Hallett, 2007). Beyond its cortical applications, TMS can modulate other CNS structures remotely, including spinal cord circuitry (Nardone et al., 2015; Anderson et al., 2022). Several studies show that it activates corticospinal neurons primarily through transsynaptic mechanisms (Ridding & Rothwell, 2007; Siebner et al., 2022; Di Lazzaro & Rothwell, 2014; Tazoe & Perez, 2015). Particularly, when stimulating the right M1-HAND representation targeting the left First Dorsal Interosseous (FDI) muscles, the main descending pathway is the left lateral corticospinal tract, which terminates on alpha motoneurons in the left ventral horn of the lower cervical cord (Kimura, 2000; Chiba et al., 2015). Neuroanatomical and neurophysiological work indicates that intrinsic hand muscles, including FDI, receive their dominant innervation from segments corresponding to C8–T1, located approximately at C7– T1 vertebral levels (Kimura, 2000; Chiba et al., 2015). To capture the downstream effects of cortical stimulation in vivo, concurrent TMS with functional magnetic resonance imaging (TMS– fMRI) has emerged as a powerful approach for investigating TMS effects on brain networks, with TMS-induced BOLD responses varying with stimulation intensity and cognitive state and showing network-level effects that extend beyond the targeted cortical area (Mizutani-Tiebel et al., 2022). However, while these effects are well characterized in the brain, the downstream impact of cortical TMS on spinal circuitry remains largely unexplored in neuroimaging and has mostly been inferred from electrophysiological measures such as motor-evoked potentials or reflexes, leaving a gap in our understanding of how TMS modulates spinal circuits in vivo. To the best of our knowledge, no previous work has systematically mapped TMS-evoked activity in the human spinal cord using spinal fMRI, which offers spatially resolved access to segmental gray and white matter territories and can therefore reveal patterns that are inaccessible to indirect physiology-based measures.

In this study, we assess spinal cord responses to cortical TMS using spinal cord fMRI. TMS pulses at varying intensities are applied to the M1-HAND area of the primary motor cortex, indirectly targeting motor neuron populations in the ventral horns of the cervical cord and evoking involuntary motor responses. Based on the known corticomotoneuronal projections to intrinsic hand muscles, we therefore expected TMS-evoked spinal fMRI activation to be maximal in the left lateral corticospinal tract and left ventral horn within C7–T1, with comparatively weaker activation outside these segments. This approach enables controlled, non-invasive stimulation of the motor pathway and provides insight into the functional organization and activation patterns of spinal motor circuits, providing a non-invasive platform for investigating TMS-modulated changes in spinal circuitry in vivo and for probing the top-down regulation of spinal circuits.

## Methods

### 2.1. Participants

Twenty-two healthy participants were enrolled in the study (age: 23.5 ± 3.2 years, sex: 10 females). All participants were right-handed and had no history of neurological disorders. Four participants had to be excluded from the analysis (see 2.5). The study was conducted in accordance with the Declaration of Helsinki and was approved by the Commission Cantonale d’Ethique de la Recherche Geneve (CCER, Geneva, Switzerland, 2021-01887). All participants provided their informed consent and were screened for MRI and TMS exclusion criteria.

### 2.2. Experimental Protocol

We developed an experimental setup integrating transcranial magnetic stimulation with spinal cord fMRI (TMS–SCfMRI), extending previously validated concurrent TMS-fMRI protocols Navarro de Lara et al., 2015, 2017) (see Figure 1A for setup). Each participant completed two identical sessions (Session-1 and Session-2) one week apart, each consisting of a preparation phase followed by concurrent TMS–SCfMRI acquisition. The acquisition phase included a brief preparation period (approximately 30 minutes) followed by TMS delivery during spinal cord fMRI. During preparation, the stimulation target in the right M1-HAND area was localized outside the scanner using an MC-B70 coil (MagPro X100, MagVenture) by identifying the scalp site that produced a discernable muscle contraction in the left hand; this hotspot was marked and then targeted in the scanner with an MR-compatible MRI-B91 coil (MagPro XP, MagVenture) positioned tangentially over the mark in a posterior–anterior orientation at approximately 45◦ to the midline. Motor threshold (MT) was determined by visual observation of MEPs via EMG as the lowest intensity reliably evoking motor potential (mean MT in the scanner = 88.4% of maximum stimulator output ± 3.9 SD), following scanner-adapted MT estimation procedures and recommendations for concurrent TMS-fMRI (Navarro de Lara et al., 2015, 2017, Mizutani-Tiebel et al., 2022). To accommodate TMS within the scanner and maintain whole-brain coverage for an extended imaging protocol that included separate brain fMRI runs, two 7-channel MR receive coil arrays were used, one placed beneath the TMS coil and another over the contralateral hemisphere (Figure 1A), following the configuration described in (Navarro de Lara et al., 2017). Although these head coils were not used for the data reported here, their presence increased the distance between the TMS coil and the scalp, adding a technical constraint that likely reduced effective stimulation intensity. Single-pulse TMS was then delivered during spinal fMRI using twelve blocks of five pulses each at three intensities, with four blocks per intensity presented in pseudorandom order and separated by 25 s rest intervals (Figure 1B), and a custom MATLAB script synchronized pulse delivery to scanner triggers to minimize imaging artifacts.

**Figure 1:**
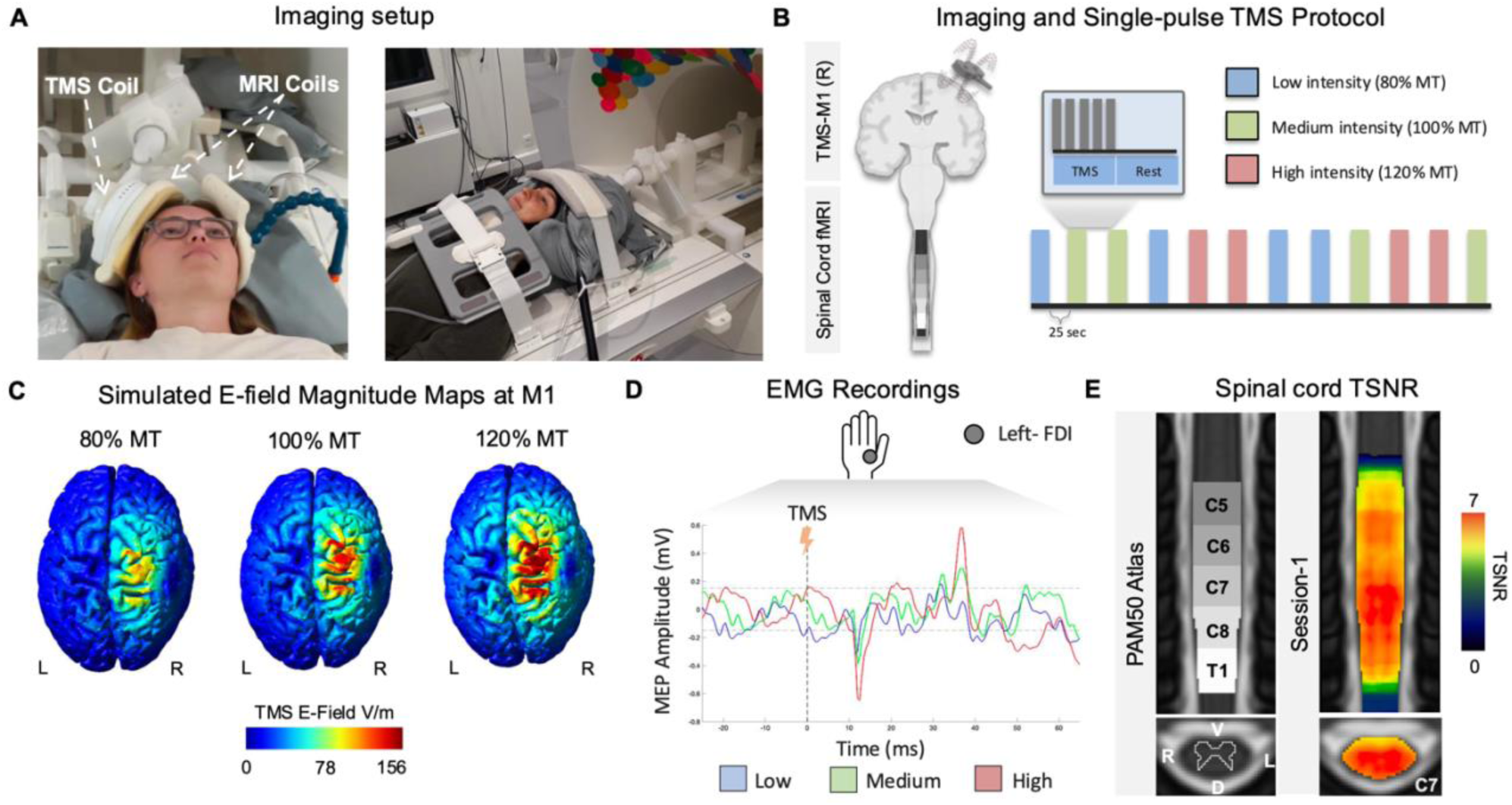
Concurrent TMS–spinal cord fMRI setup and data quality. (A) Schematic of the experimental setup showing MR-compatible TMS and MRI receiving coils and the 32-channel spinal coil array used to acquire cervical spinal cord fMRI during TMS over right M1-HAND. (B) Single-pulse TMS protocol: 12 blocks of 5 pulses each delivered at three intensities (Low 80% MT, Medium 100% MT, High 120% MT) in pseudorandom order during spinal cord fMRI. (C) Simulated electric field (E-field) distributions in a representative subject for 80%, 100%, and 120% MT, illustrating consistent targeting and increasing E-field magnitude within the precentral M1-HAND region. (D) Example EMG recording from the left FDI muscle in the same subject, showing MEPs evoked at the three TMS intensities relative to the setup-specific baseline noise band (± 0.15 mV) and the TMS pulse at 0 ms (E) Group-level temporal SNR (tSNR) map of the cervical spinal cord overlaid on the PAM50 atlas, demonstrating adequate signal quality and coverage across lower cervical segments, including C7–T1. The photograph in panel A depicts one of the authors demonstrating the experimental setup.

### 2.3. Transcranial Magnetic Stimulation and Electromyographic Recordings

TMS was administered using a MagPro X100 stimulator with an MR-compatible MRi-B91 figure-of-eight coil (Magventure, Farum, Denmark), targeting the right M1-hand area representing the left FDI muscles. Single-pulse TMS was delivered at three intensity levels: 80% MT (Low), 100% MT (Med), and 120% MT (High) (Figure 1B). Surface EMG was recorded from the left FDI, and the stimulation hotspot was defined as the scalp location eliciting the largest and most consistent motor-evoked potentials (MEPs) in FDI (Figure 1D). Peak-to-peak MEP amplitudes were computed from the raw EMG within a 50 ms window after each TMS pulse, with trials containing significant noise artefacts excluded after visual inspection; low-frequency 50 Hz noise from the MR environment did not affect MEP estimation, and radiofrequency interference was avoided by pausing image acquisition for 1430 ms around each pulse.

### 2.4. Spinal Cord Magnetic Resonance Imaging

Imaging data were acquired on a 3.0-T Siemens Prisma scanner at the Human Neuroscience Platform (Campus Biotech, Geneva, Switzerland) using a 32-channel spine coil together with a 16-channel body coil. Functional images of the cervical spinal cord were obtained with a ZOOMit gradient-echo EPI sequence (TR = 2.5 s, TE = 34 ms, flip angle = 80°, FOV = 48 x 144 mm^2^, in-plane resolution = 1 x 1 mm^2^, slice thickness = 3 mm), with 28 axial slices covering C2–T1 and spinal-focused shimming performed before acquisition (Kinany et al., 2019). Each run contained 293 volumes (12 min 20 s). A high-resolution T2-weighted anatomical image of the cervical cord was acquired using a 3D SPACE sequence (TR = 1660 ms, TE = 133 ms, echo train length = 73, flip angle = 140°, resolution = 0.4 x 0.4 x 0.8 mm^3^, sagittal orientation). Cardiac and respiratory signals were recorded with a pulse oximeter and a respiratory belt for subsequent physiological noise modeling. TMS pulses were delivered concurrently within the spinal fMRI runs on every second TR with a 200 ms post-trigger delay to avoid interference with MR acquisition, using an in-house MATLAB script synchronized to the scanner triggers.

### 2.5. Spinal Cord fMRI Preprocessing

All spinal cord images were converted to NIFTI and preprocessed with in-house Python pipelines using FSL, the Spinal Cord Toolbox (De Leener et al., 2017), and Nilearn, following standard spinal cord fMRI preprocessing procedures described in the literature (Eippert et al., 2017; Kinany et al., 2023). Preprocessing comprised: (1) slice-wise motion correction within a cylindrical mask around the cord, with framewise displacement (FD) computed at each level and mean spinal cord FD of 0.09 ± 0.04 mm in Session-1 and 0.11 ± 0.05 mm in Session-2 (Figure S3), and no run exceeding a 0.3 mm threshold; (2) automatic segmentation of spinal cord and cerebrospinal fluid on anatomical and mean functional images using SCT deep-learning tools, followed by visual inspection and manual correction where necessary; (3) physiological denoising using a RETROICOR-style model with slicespecific cardiac, respiratory, interaction, and CSF regressors (32 physiological regressors plus one CSF regressor) to reduce physiological artifacts; (4) spatial normalization of functional data to the PAM50 template (De Leener et al., 2018) via intermediate anatomical-to-template and functional-to-anatomical registrations, with the resulting transforms concatenated into a single functional-to-template warp; and (5) spatial smoothing of the denoised, normalized images with a 3D Gaussian kernel of 2 x 2 x 6 mm^3^ full width at half maximum. Following slice-wise motion correction, we computed temporal signal-to-noise ratio (tSNR) on minimally processed functional images by dividing, for each voxel, the mean signal intensity across time by its standard deviation, providing a voxel-wise measure of spinal cord signal stability over the run. Temporal SNR in C5–T1 segments averaged 6.04 ± 0.30 in Session-1 and 5.92 ± 0.42 in Session-2 (Figure 1E and Figure S3), indicating stable spinal signal quality during concurrent TMS–SCfMRI; datasets with excessive residual motion or poor registration were excluded, yielding a final sample of nineteen participants for Session-1 and twenty-one participants for Session-2. To derive temporally sharpened representations of neural activity from these preprocessed time series, we next applied the Total Activation framework based on regularized hemodynamic deconvolution (Karahanoğlu & Van De Ville, 2015; Karahanoğlu et al., 2013). Specifically, TA was used to transform the motion-corrected, denoised, normalized, and smoothed spinal cord fMRI data into activity-inducing time courses for all subsequent analyses.

### 2.6. Subject Categorization Based on TMS–EMG Output

Inter- and intra-individual variability in physiological responses to non-invasive brain stimulation is well documented in TMS studies (Pellegrini et al., 2018a, 2018b; Ammann et al., 2017; Wiethoff et al., 2014; Nuzum et al., 2016). Such variability is often quantified using input-output (IO) relationships between stimulation intensity and MEP amplitude, with steeper curves interpreted as greater recruitment of corticospinal pathways and increased corticospinal excitability (Pellegrini et al., 2018c; Vetter et al., 2023). In this work, our central hypothesis was that spinal BOLD responses should depend on how effectively TMS engages the descending motor system. However, considering the technical challenges inherent to concurrent TMS-SCfMRI we expected the IO curves can potentially be degraded by coil positioning, minor drifts, and EMG setup rather than intrinsic physiology. Thus, our first step in each session was to characterize corticospinal recruitment in each subject using individual IO curves and to verify that TMS pulses produced a reliable EMG response before interpreting spinal activation. For every subject and session, we computed peak-to-peak MEP amplitudes for each TMS pulse, averaged within blocks of five pulses, and then across blocks for Low (80% MT), Medium (100% MT), and High (120% MT) intensities. We examined these IO profiles relative to a setup-specific EMG noise floor (± 0.15 mV) estimated from baseline periods. To operationalize the distinction between clear and ambiguous IO behavior, we applied two subject-level criteria: (i) the mean peak-to-peak MEP amplitude at Low, Medium, and High intensities had to be at least 0.20 mV, ensuring responses were robustly supranoise in our setup, and (ii) mean MEP amplitudes had to increase across the three tested intensities with a minimum positive change of 0.15 mV at each step, reflecting a clear intensity-dependent gain in corticospinal output. Applying these operational thresholds, we classified subjects as High Output Group (HOG) when both criteria were satisfied and Low Output Group (LOG) otherwise, yielding 11 HOG and 8 LOG subjects in Session-1 and 14 HOG and 5 LOG subjects in Session-2; this provided an empirically grounded, hypothesis-consistent stratification of corticospinal output profiles for subsequent spinal fMRI analyses.

To ensure that this grouping did not simply reflect gross targeting errors or hardware artifacts, we complemented the EMG-based inspection with individualized E–field simulations over right M1-HAND. These simulations confirmed broadly equivalent M1-HAND field coverage and strength across subjects in both sub groups, at 80–120% MT (Figure S1), controlling against possbile systematic differences in coil positioning or stimulator performance as the primary driver of the heterogeneity in MEP output, although residual variability is expected in such a constrained setup. At 120% MT, where clear TMS-evoked responses are expected, only HOG subjects showed MEP amplitudes that reliably exceeded the EMG noise floor, whereas LOG subjects tended to remain near baseline, consistent with the representative example in Figure S2. Importantly, we treat this grouping as an operational stratification of effective recruitment of corticospinal system under known constraints of concurrent TMS–SCfMRI, not as a neurophysiological subtype, and we use it to focus mechanistic interpretations on subjects where the peripheral motor response behaves as experimentally expected while still acknowledging that spinal BOLD responses can be detected more broadly. This approach builds conceptually on prior work using IO curves to quantify responsiveness in TMS studies, but explicitly adapts the classification to the technical limitations and data-quality considerations inherent in spinal fMRI.

### 2.7. fMRI Data Analysis

Spinal cord fMRI data were analyzed voxel-wise within a General Linear Model (GLM) framework using FSL and Nilearn, applied to the preprocessed activity-inducing time series obtained from the TA step. First-level models were estimated for each subject and session with task-related regressors and motion parameters as confounds, without additional HRF convolution. Subject-level contrast and Z-statistic maps were then entered into non-parametric group analyses using FSL randomise with Threshold-Free Cluster Enhancement (TFCE), and 10,000 permutations to account for the spinal cord’s small cross-section and non-Gaussian noise characteristics (Nichols & Holmes, 2002; Winkler et al., 2014; Worsley et al., 1992; Frostell et al., 2016) and inference constrained to C5–T1 spinal levels. From the resulting group-level statistical maps, we quantified (i) the percentage of active voxels within C5–T1 spinal levels, using a group-level FWE-corrected threshold of *p* < 0.005 and (ii) the spatial overlap between significant clusters and PAM50 atlas-based white-matter tracts using Dice coefficients. We also extracted parameter estimates from voxels within gray-matter masks that survived group-level FWE-correction at *p* < 0.005 and used these values for post-hoc paired *t*-tests between spinal levels. Three GLM specifications were used: (A1) a block-based model with a single regressor capturing all TMS events to test for TMS-evoked versus baseline BOLD responses; (A2) a block-based model with separate regressors for Low (80% MT), Medium (100% MT), and High (120% MT) intensities to assess intensity-dependent modulation; and (A3) an event-related model incorporating trial-wise peak-to-peak MEP amplitudes from EMG recordings (Section 2.3), with one unmodulated regressor (constant MEP value) and one parametric modulator (trial-wise MEP amplitude) orthogonalized with respect to the first to avoid imposing a one-to-one mapping between zero MEP and zero BOLD response (Mumford et al., 2015).

## Results

### 3.1. TMS-Induced MEP Output and Spinal Activation

We first characterized TMS–EMG output by examining peak-to-peak MEP amplitudes at Low (80% MT), Medium (100% MT), and High (120% MT) intensities, and stratified subjects into HOG and LOG using the a priori, subject-level supranoise and intensity-dependent IO criteria described in Section 2.6. Building on this subject-level stratification, Figure 2A provides a complementary group-level view of the resulting IO profiles: as expected from the definition, HOG subjects show larger MEP amplitudes that increase with intensity and remain clearly above the EMG noise floor, whereas LOG subjects display flatter IO curves with amplitudes closer to baseline. Beyond simply mirroring the subject-wise criteria, Figure 2A demonstrates that these patterns also hold at the group level, i.e. showing significant differences across all three intensities within HOG (paired tests, *p* < 0.05) and systematic HOG–LOG differences at each intensity (between-group tests, *p* < 0.05) in both sessions, thereby confirming that the subject-level stratification yields two internally consistent and clearly separable IO profiles that provide a meaningful basis for interpreting downstream spinal fMRI effects.

**Figure 2:**
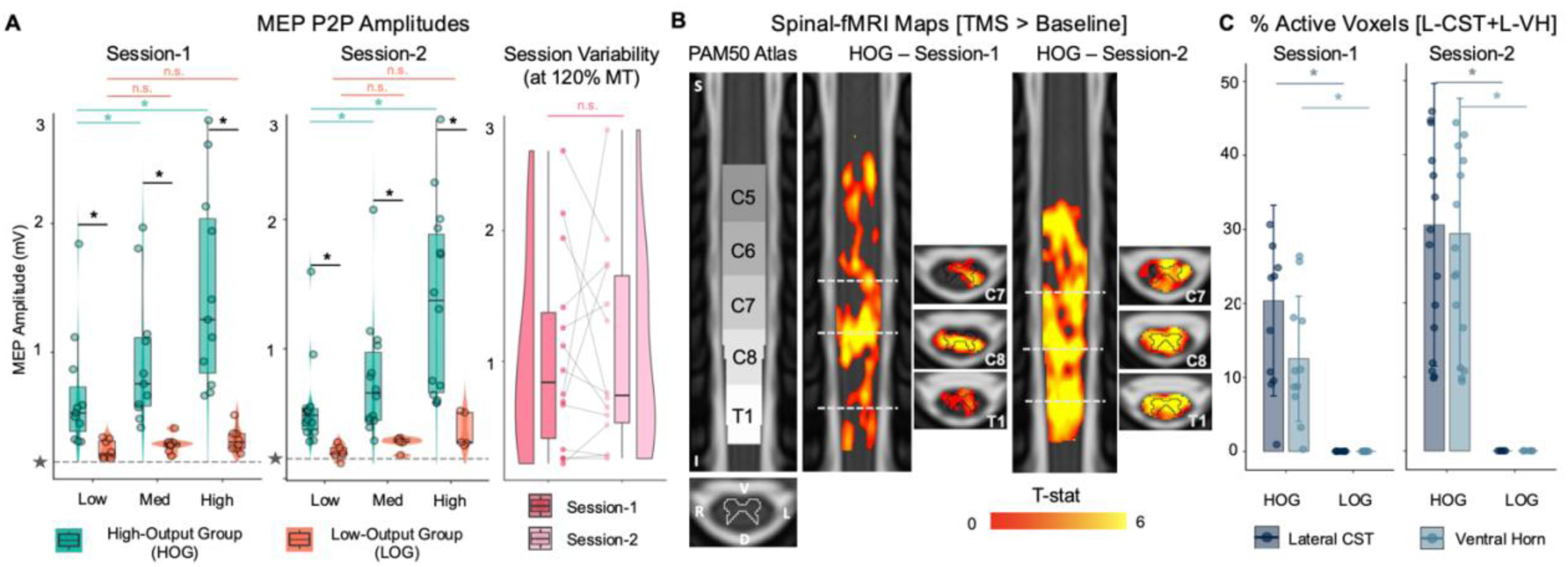
TMS–EMG output and spinal activation in High Output Group (HOG) vs. Low Output Group (LOG). (A) Peak-to-peak motor-evoked potential (MEP) amplitudes across three TMS intensities (Low 80% MT, Medium 100% MT, High 120% MT) for subjects classified as HOG and LOG in Session-1 (*n* = 11 HOG, *n* = 8 LOG) and Session-2 (*n* = 14 HOG, *n* = 5 LOG) based on input-output profiles relative to the EMG noise floor (Section 2.6). Within-subject intensity effects in HOG are assessed with paired t-tests (*: *p* < 0.05, n.s.: otherwise), and between-group differences between HOG and LOG at each intensity are assessed using a Welch’s t-test (*: *p* < 0.05, n.s.: otherwise). An additional paired box plot compares mean peak-to-peak MEP amplitudes at 120% MT between the common set of subjects present in both sessions, illustrating session-level variability in high-intensity responses. (B) Group-level spinal cord activation maps from the A1 model (main effect of TMS > baseline), showing t-statistic values within clusters surviving FWE-corrected *p* < 0.005 (0 ≤ *t* ≤ 6) across C5–T1 in HOG for Session-1 and Session-2, displayed on the PAM50 template (axial and coronal views). The dotted lines on coronal views mark the represented axial location. S = superior, I = inferior, L = left, R = right, D = dorsal, V = ventral. No significant activation was observed in LOG at the same threshold in either session. (C) Percent of significantly active voxels within PAM50 masks of the left lateral corticospinal tract and left ventral horn gray matter at C7–T1 (*p* < 0.005, FWE-corrected) for HOG and LOG in Session-1 and Session-2.; HOG-LOG differences are assessed using two-sample tests (*p* < 0.05).

To further assess consistency of TMS–EMG output across sessions, we additionally compared mean peak-to-peak MEP amplitudes at 120% MT between Session-1 and Session-2 in the subset of subjects present in both sessions (Figure 1A, right). Despite identical targeting procedures and stimulation parameters, individual high-intensity MEP amplitudes varied noticeably between sessions, as reflected by paired scatter points and connecting lines. Across the subjects, test–retest reliability was only modest (Pearson *r* = 0.43, *p* = 0.12; ICC(A,1) = 0.45, 95% CI: [−0.11 0.79], *p* = 0.053), indicating poor–to–moderate agreement with considerable uncertainty. This pattern suggests that session-level differences in effective recruitment and/or setup constraints can markedly influence peripheral output, motivating a session-wise analysis of spinal fMRI rather than pooling across sessions, and reinforcing our choice to base mechanistic inferences on HOG/LOG stratification within each session.

Building on this grouping, we then asked whether spinal activation reflects these differences in peripheral output. Using the A1 model (single regressor capturing all TMS events) and a strict threshold (*p* < 0.005, FWE-corrected) applied uniformly to HOG and LOG, group-level spinal maps across C5-T1 revealed robust TMS-evoked activation only in HOG subjects in both sessions, with clusters spanning cortiscospinal tracts, dorsal columns, intermediate zones, and ventral horns, and extending rostrocaudally from approximately C5 to T1 (Figure 2B). No voxels survived the same threshold in LOG for either session, and thus LOG maps are not displayed. To test whether activation in HOG aligns with the expected neuroanatomical pattern for right M1-HAND stimulation and to build further confidence that HOG reflects reliable recruitment of the hypothesized spinal territories rather than improper stimulation or sensory artefacts, we quantified the percentage of active voxels within PAM50 masks of the left lateral corticospinal tract and left ventral horn gray matter at C7–T1, applied to the significant clusters in Figure 2B. As shown in Figure 2C, HOG subjects in both sessions exhibited non-zero fractions of active voxels in these masks, whereas LOG subjects showed 0% active voxels at the same threshold, and HOG–LOG differences in percentage of active voxels were significant for both tracts and both sessions (*p* < 0.05). Together, these results show that our HOG/LOG stratification is justified: only HOG subjects, who exhibit the experimentally expected supranoise IO behavior, also show robust spinal fMRI activation in the anatomically predicted left lateral CST and ventral horn at C7–T1, whereas LOG subjects do not exhibit detectable activation in these territories under conservative thresholds. This supports using HOG as the primary subgroup for mechanistic spinal analyses while acknowledging the technical constraints of concurrent TMS–SCfMRI.

### 3.2. TMS-Intensity Dependent Effects on Spinal Cord BOLD Signal

Having established that TMS–EMG output separates subjects into High Output and Low Output Groups and that only HOG shows robust, anatomically plausible spinal activation (Section 3.1), we next tested whether spinal cord BOLD responses varied systematically with TMS intensity within HOG, focusing on Session-2 where the number of HOG subjects was highest and IO behavior most clearly expressed (Session-1 HOG results are reported in Figure S4). Using the A2 model (separate regressors for Low, Medium, and High intensities; Section 2.7), and based on the MEP findings, we hypothesized a progressive increase in both the extent and neuroanatomical spread of activation from Low to High, with predominantly dorsal (sensory) activation at Low, mixed dorsal–ventral involvement at Medium, and dominant contralateral ventral (motor) activation at High. Group-level maps confirmed a clear intensity-dependent increase in the proportion of active spinal voxels (FWE-corrected *p* < 0.005; Figure 3), with mean active-voxel percentages of 5.26 ± 3.37 for Low, 28.84 ± 8.78 for Medium, and 48.81 ± 19.74 for High, indicating a graded increase in activation with stimulation intensity. Spatially, Low intensity produced limited activation mainly in the dorsal column at C7–C8, the contralateral dorsal horn at C7, and the descending corticospinal tract at T1, whereas Medium intensity recruited both dorsal and ventral horns, with dominant contralateral dorso–ventral activation from C7 to T1, ipsilateral activation at C5, and clusters in the contralateral ventral horn and descending tract at C6. High intensity yielded the most extensive pattern, with activation encompassing the intermediate zone and bilateral ventral horns from C6 to T1, additional clusters in the ipsilateral corticospinal tract (C7–T1), and dorsal column activity spanning C5–T1. Parameter estimates extracted across spinal levels (Section 2.7) increased significantly from Low to High at all levels (*p* < 0.05), with C6 and T1 also showing significant Medium–High differences, demonstrating a robust intensity-dependent increase in spinal BOLD responses in Session-2. Session-1 also revealed a similar intensity-dependent effect, however, comparatively weaker than Session-2 (Figure S4).

**Figure 3:**
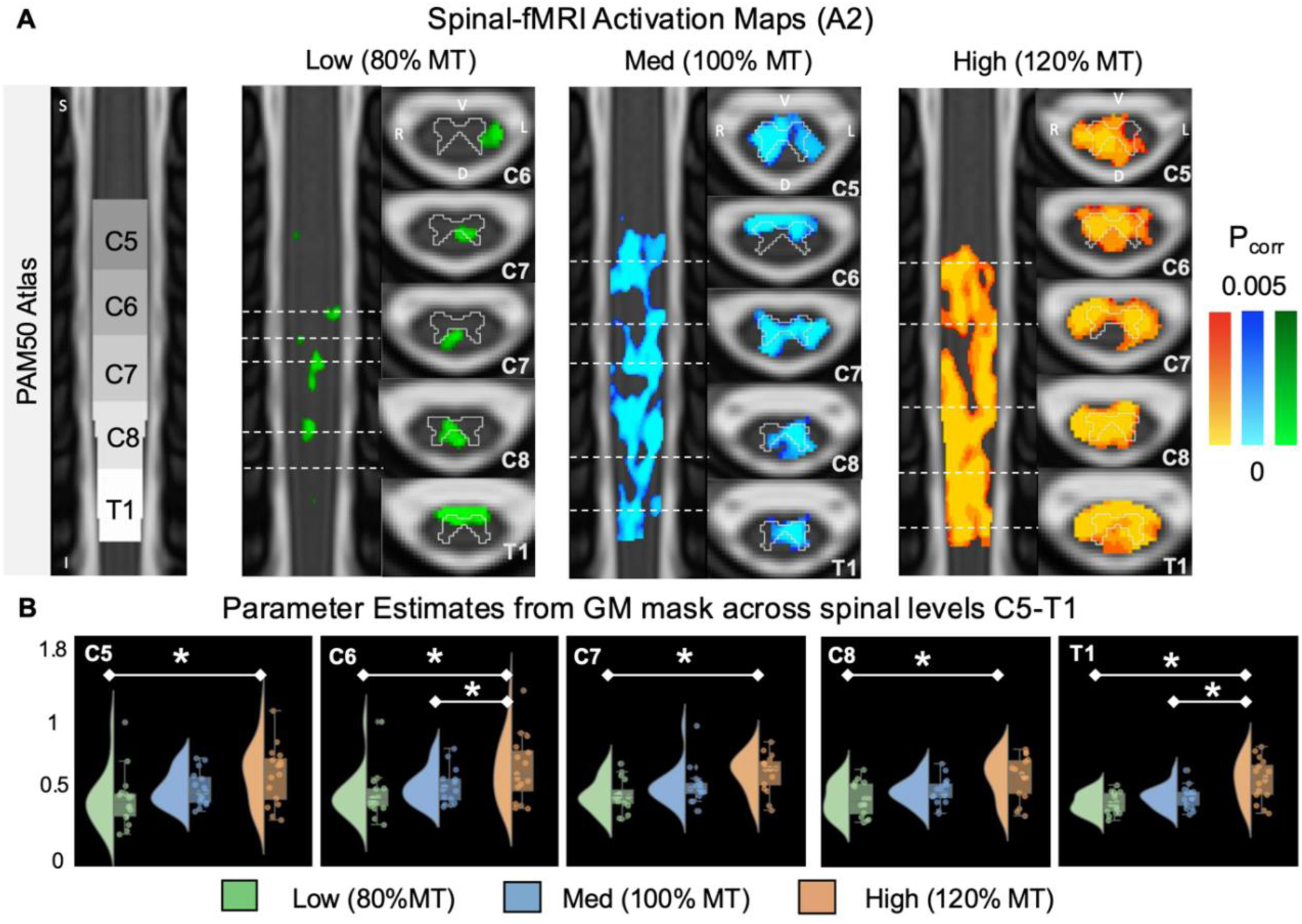
Intensity-Dependent Spinal BOLD Responses to TMS. (A) Group-level spinal activation maps for three TMS intensities (Low, Medium, High), thresholded at *p* < 0.005 (FWE-corrected). Axial slices are overlaid on the PAM50 template and labeled by spinal level (C5–T1). The dotted line on coronal views marks the represented axial location. S = superior, I = inferior, L = left, R = right, D = dorsal, V = ventral. (B) Parameter estimates from activated voxels within the gray matter across spinal levels (C5–T1) for Low, Medium, and High TMS intensities, illustrating intensity-specific BOLD response amplitudes across spinal levels.

**Figure 4:**
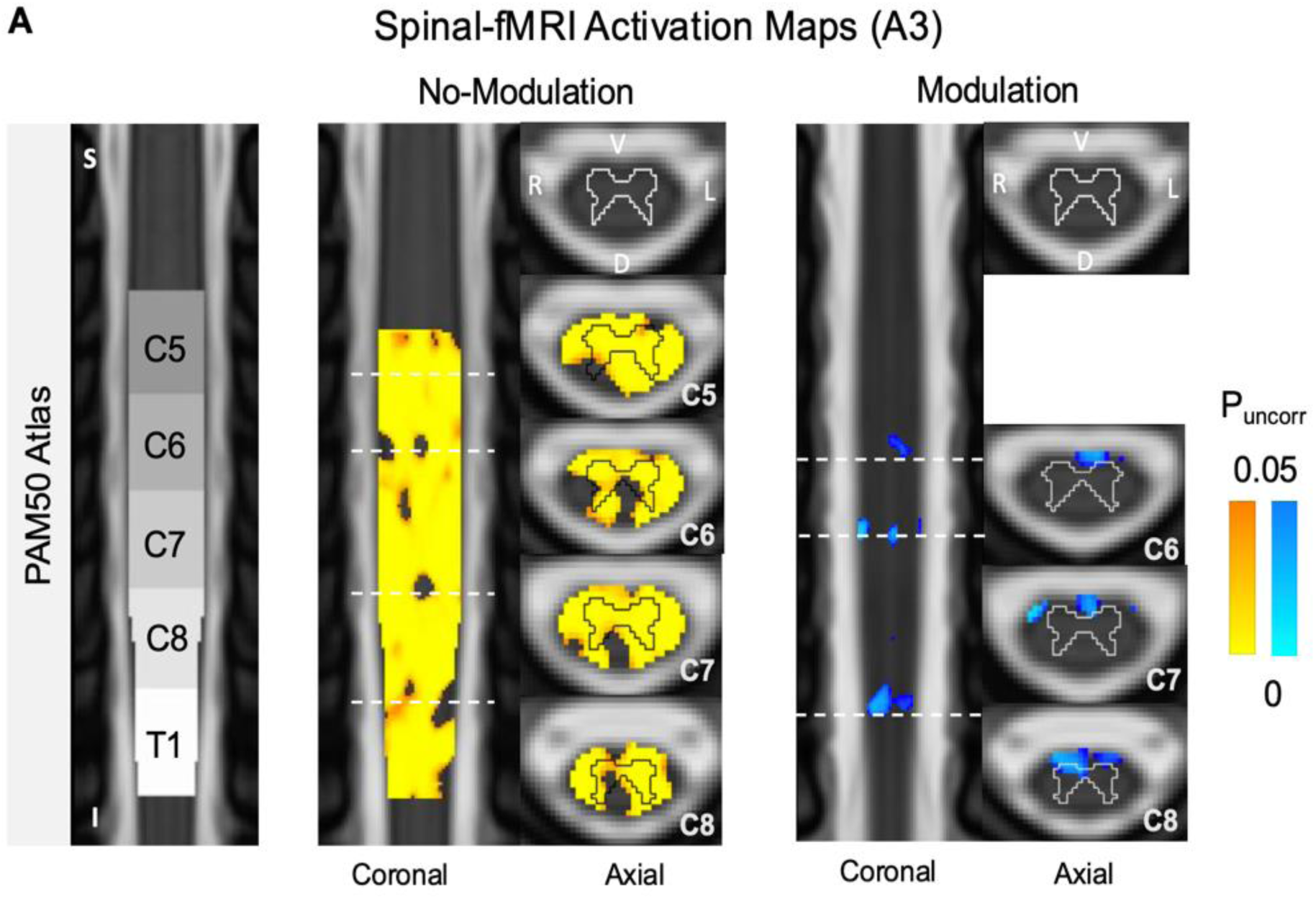
Trial-by-trial Modulation of Spinal BOLD Signal by MEP Amplitudes. Group-level activation map for the unmodulated regressor (reflecting trial-wise TMS events) shows distributed activation spanning C5–T1, across both gray and white matter regions, thresholded at *p* < 0.05, FWE-uncorrected. The dotted line on coronal views marks the represented axial location. S = superior, I = inferior, L = left, R = right, D = dorsal, V = ventral.

### 3.3. Effect of Modulation on Spinal Cord BOLD Signal

To test whether trial-by-trial variations in MEP amplitude were associated with corresponding fluctuations in spinal BOLD responses, we analyzed activation maps from the MEP-based parametric GLM (A3; Section 2.7) in HOG of Session-2, which included an unmodulated regressor capturing the presence of TMS events and a parametric regressor whose amplitude scaled with the evoked MEPs. Group-level maps estimated with non-parametric permutation testing, and TFCE showed that, at an exploratory threshold of *p* < 0.05 (FWE-uncorrected), the unmodulated regressor produced activation spanning C5–T1 with distributed involvement of gray and white matter regions (Figure 4), consistent with the block-based TMS > baseline contrast in A1. By contrast, the MEP-modulated regressor yielded activation (also at *p* < 0.05, FWE-uncorrected) primarily confined to the descending corticospinal tract, with peak overlap at C6 (Dice coefficient 0.34 ± 0.14), C7 (0.51 ± 0.15), and C8 (0.44 ± 0.13), and no detectable activation in the ventral horns, aligning with the hypothesis that modulation is strongest in descending motor pathways rather than in ascending sensory tracts. Comparison of descending corticospinal tracts and ventral horn regions (DT) versus ascending tracts (AT) confirmed this specificity: Dice coefficients were significantly larger in descending than ascending pathways (*DC_AT_* = 0.0 vs. *DC_DT_* = 0.11 ± 0.05, *p* < 0.0001, Mann–Whitney U test across non-zero slices; Figure 4), indicating that trial-by-trial peripheral motor output provides a complementary, pathway-specific marker of spinal BOLD modulation beyond stimulation intensity alone. Session-1 also revealed a similar effect, however, comparatively stronger than Session-2 (Figure S5).

**Figure 4:**
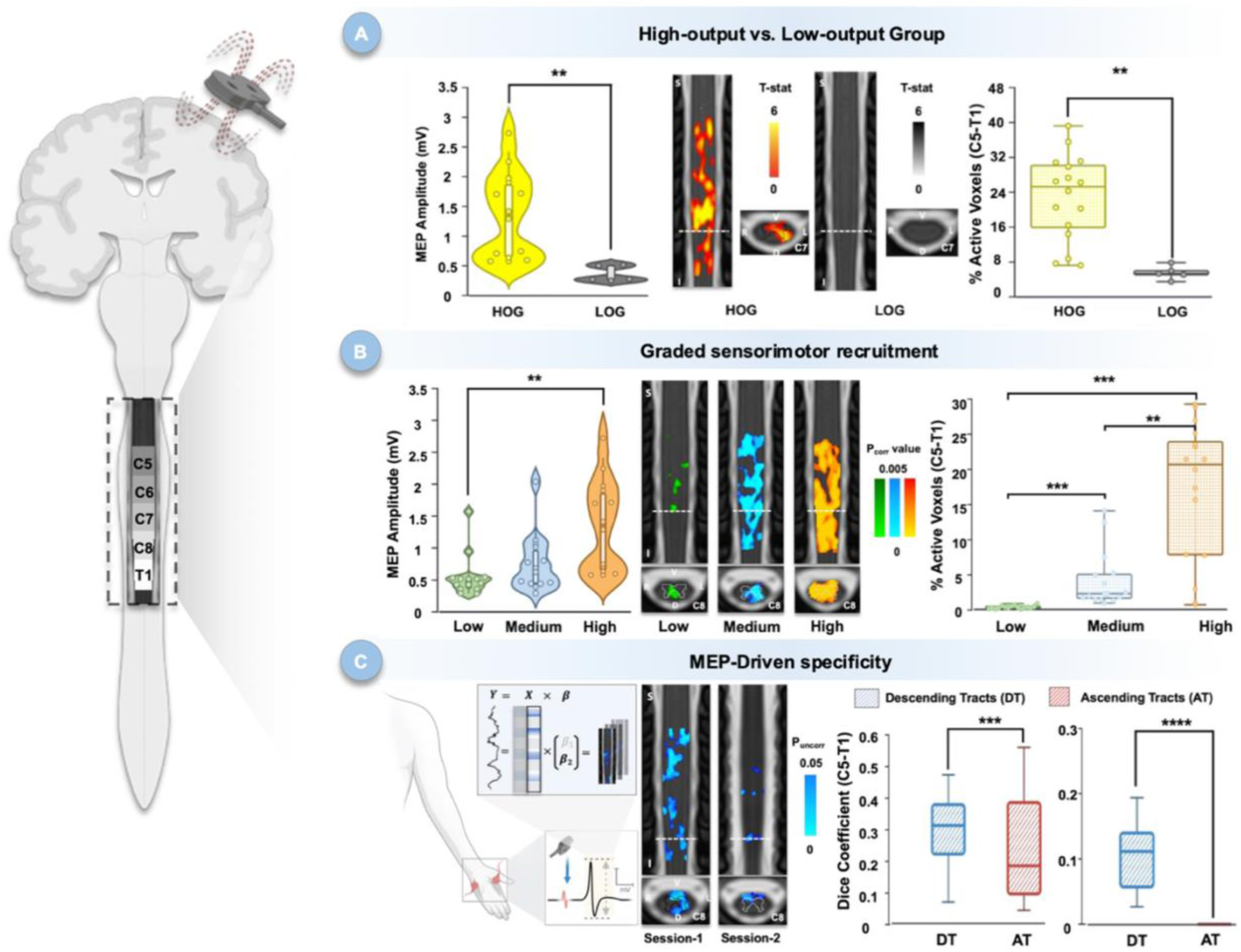
TMS-evoked spinal fMRI reveals motor output–dependent, intensity-driven, and pathway-specific recruitment of human spinal circuits. A. HOG vs. LOG. Left: Participants classified by motor output responsiveness across TMS intensities. Middle: Spinal fMRI reveals widespread activation in HOG compared to sparse activation in LOG. Right: HOG show a higher percentage of active spinal voxels than LOG. B. Graded Sensorimotor Recruitment. Left: In HOG, MEP amplitudes progressively increase from low to high TMS intensities. Middle: Spinal fMRI maps show expanding spatial extent of activation with increasing intensity, involving dorsal and ventral horns and descending tracts. Right: Percentage of active spinal voxels rises significantly from low to high intensity conditions. C. MEP-Driven specificity. Left: Schematic of trial-wise MEP-based parametric modulation, showing EMG acquisition, MEP extraction, and integration into the GLM design matrix. Middle: Modulated regressor highlights stimulation-relevant regions, including descending corticospinal tracts and ventral horns, with broader caudal coverage in Session-1. Right: Descending tracts show significantly greater activation than ascending tracts across sessions, confirming motor-pathway specificity to stimulation.

## Discussion

In this study, we used transcranial magnetic stimulation of the motor cortex combined with spinal cord fMRI and a multimodal analysis framework to examine how cortical stimulation propagates through descending motor pathways and interacts with spinal circuits. For decades, TMS studies have quantified these effects indirectly using measures such as EMG, MEPs, or spinal reflexes, which are sensitive to corticospinal output but provide limited information about the spatial distribution of activity within the cord and make it difficult to separate spinal from supraspinal contributors. By providing a non-invasive, spatially resolved window onto spinal BOLD responses during cortical neuromodulation, spinal fMRI complements these traditional readouts and, as implemented in our framework, allows in vivo mapping of how cortical stimulation engages specific spinal segments and pathways in humans.

### Differential Spinal Activation in High Output and Low Output Groups

The first analysis (A1) showed that TMS over M1 evokes reliable BOLD responses in the cervical spinal cord, engaging gray and white matter regions including ventral horns, dorsal columns, and intermediate zones, but with marked differences between High Output Group (HOG) and Low Output Group (LOG) subjects (Figure 4A). In HOG, who exhibited the expected increase in MEP amplitudes with higher TMS intensities, robust activation patterns were observed across sensorimotor spinal regions, consistent with effective recruitment of descending corticospinal volleys. This interpretation aligns with electrophysiological work showing that TMS over M1 activates fast-conducting corticospinal fibers and elicits D- and I-waves arising from monosynaptic activation of pyramidal tract neurons whose volleys propagate to spinal motoneurons (Di Lazzaro et al., 2010; Di Lazzaro & Rothwell, 2014) and interneurons, particularly within the propriospinal system of the cervical cord (Di Lazzaro et al., 1998; Darling et al., 2006). In contrast, LOG showed no detectable recruitment of the hypothesized lateral CST and ventral horn territories at C7–T1 under conservative thresholds, consistent with more limited effective TMS–EMG coupling in the constrained scanner environment. As highlighted by Darling et al. even at fixed TMS intensities MEP amplitudes can vary substantially due to fluctuations in cortical and spinal excitability, propriospinal contributions, and cognitive state, and these sources of variability are likely amplified in an innovative multimodal setup such as concurrent TMS–SCfMRI, where coil positioning, minor drifts, EMG configuration, and subject compliance further influence the efficacy of stimulation (Darling et al., 2006; Carroll et al., 2001). In this context, spinal BOLD responses reflect not only the nominal stimulation intensity but also the effectiveness with which descending pathways are recruited and translated into peripheral motor output, reinforcing the need for an operational HOG/LOG stratification that explicitly acknowledges both physiological and technical contributors. Our findings extend this literature by demonstrating that such variability in descending recruitment can be visualized and quantified with spinal fMRI at the level of specific segments and pathways, particularly the left lateral CST and ventral horn at C7–T1 in HOG, providing a non-invasive readout of sensorimotor pathway function that explicitly incorporates both physiological and setup-related sources of variability.

### Intensity-Dependent Spinal Recruitment Reveals Graded Sensorimotor Engagement

In the second analysis (A2), we observed a progressive increase in the extent and spatial distribution of spinal BOLD activation as TMS intensity increased, with higher intensities recruiting a larger proportion of spinal voxels and broader rostrocaudal territories (Figure 4B). This graded behavior is consistent with other neurophysiological work such as EMG recruitment curves showing a sigmoidal increase in MEP amplitude once stimulation exceeds MT (Carroll et al., 2001; Kukke et al., 2014; Koponen et al., 2024; Krile et al., 2023), TMS– EEG studies linking larger TMS-evoked potentials in motor cortex to higher MEP amplitudes and stronger descending output, and in clinical populations with partially preserved corticospinal pathways (Neva et al., 2016; Nardone et al., 2015; Chowdhury et al., 2023). Although MEP amplitude has traditionally served as an indirect measure of corticospinal excitability (Di Lazzaro & Rothwell, 2014), its relationship to spinal BOLD has only recently become accessible via TMS–spinal fMRI. Given established neurovascular coupling and the known organization of descending motor pathways increased corticospinal drive at higher TMS intensities is likely to recruit more extensive synaptic activity in spinal gray matter, particularly in ventral horn regions, thereby elevating local metabolic demand and generating stronger hemodynamic responses detectable with spinal fMRI (Petrichella et al., 2017, Desmons et al., 2023). The ability to elicit distal motor responses by stimulating the contralateral M1-HAND region provides direct evidence for antegrade, transsynaptic propagation along pre-existing corticospinal pathways (Siebner et al., 2022), enabling local cortical stimulation to drive remote activation of spinal interneurons and motoneurons. In our data, spinal activation shifted from predominantly dorsal regions at lower intensities to more pronounced ventral horn and descending tract engagement at higher intensities, mirroring cortical TMS–fMRI findings in which subthreshold stimulation preferentially enhances activity in premotor and sensory cortices, whereas suprathreshold stimulation more strongly recruits primary motor areas and produces larger corticospinal responses (Bestmann et al., 2003, Shitara et al., 2013, Mizutani-Tiebel et al., 2022). Taken together, the intensity-dependent scaling and dorsal-to-ventral progression of spinal BOLD responses in our study are consistent with graded recruitment of descending volleys and their integration within spinal sensorimotor circuits.

### Trial-Wise MEP-Based Modulation Provides Functional Specificity

Building on the robust, intensity-dependent spinal BOLD responses observed in A1 and A2, the third analysis (A3) used an electrophysiologically informed model in which trial-wise fluctuations in motor output, indexed by MEP amplitude, were used to capture known variability in TMS-elicited responses across trials (Darling et al., 2006). When TMS pulses were modeled with an unmodulated regressor, spinal activation closely resembled the A1 pattern, extending along the rostrocaudal axis and involving both gray and white matter, indicating that the spinal cord is broadly sensitive to the presence of stimulation. In contrast, the MEP-modulated regressor revealed a more selective pattern, with activation primarily localized to descending pyramidal (corticospinal) and extrapyramidal (reticulospinal, vestibulospinal) tracts that convey motor commands from the brain (Figure 4C), consistent with evidence that cortical TMS influences spinal circuitry via both pathways (Di Lazzaro et al., 2010; Fisher et al., 2012; Mooney et al., 2023). Although we expected modulation-related activation in gray matter, particularly in contralateral ventral horn and intermediate zones where descending commands are integrated, no significant clusters were detected there, potentially reflecting high trial-wise variability in motoneuron recruitment and reduced group-level effect sizes (Darling et al., 2006). Together, these results show that peripherally informed modeling adds functional specificity: the unmodulated regressor captures general TMS sensitivity of the cord, whereas the MEP-modulated regressor isolates descending motor pathways, indicating that trial-wise spinal BOLD signals can index meaningful recruitment of motor-specific circuitry beyond what is accessible with conventional block-based or unmodulated event-related designs.

By showing that cortical TMS evokes graded, intensity-dependent spinal activity in healthy individuals, our work underscores the spinal cord’s active role in sensorimotor control and paves the way for clinically meaningful spinal imaging readouts. Non-invasive stimulation approaches, such as repetitive TMS, have already shown therapeutic benefits in several neurological and psychiatric conditions, including movement disorders, where motor cortex stimulation can enhance corticospinal excitability and support recovery (Ridding and Rothwell, 2007, Lefaucheur et al., 2014; Hernandez-Navarro et al., 2025). Yet, most interventions lack spatially resolved imaging evidence linking cortical stimulation to spinal mechanisms. In this context, combining spinal fMRI with TMS and peripheral measures, as in our framework, could enable non-invasive mapping of spinal neural dynamics before and after intervention, help identify residual functional pathways, and yield individualized biomarkers of spinal circuit integrity in clinical disorders (Grefkes & Fink, 2014; Drysdale et al., 2017; Cocchi & Zalesky, 2018). Such biomarkers may guide personalized neuromodulation and spinal stimulation strategies, inform the design and optimization of brain–spine interfaces, and support longitudinal tracking of spinal plasticity, recovery trajectories, and treatment responsiveness.

At the same time, limitations of this TMS–spinal fMRI approach should be acknowledged. We observed substantial inter-individual and inter-session variability in both MEPs and spinal BOLD responses, consistent with the known variability of TMS effects across individuals and sessions (Carroll et al., 2001; Darling et al., 2006; Rafiei & Rahnev, 2022). A more systematic characterization of this variability will require accounting for differences in corticospinal excitability and anatomical-pathway organization as well as session-specific technical factors such as coil positioning, minor drifts, and EMG setup that can degrade IO curves and effective stimulation in a concurrent TMS–SCfMRI environment. Moreover, TMS-evoked BOLD responses are often weak or inconsistent at the cortical level due to factors such as coil positioning, limited spatial precision, and complex neurovascular coupling (Rafiei & Rahnev, 2022), challenges that may be at least as pronounced in spinal fMRI given additional anatomical and technical constraints. An additional limitation is that we did not continuously monitor or control low-level background EMG activity in the target muscle; small changes in tonic motoneuron drive and associated sensory afference can modulate MEP amplitudes and may therefore have contributed to some of the observed variability in peripheral output and spinal BOLD. Future studies should therefore incorporate more precise TMS targeting based on measured coil position, improved spinal fMRI acquisition schemes, continuous EMG monitoring with stricter criteria for muscle relaxation, larger cohorts, and appropriate control conditions (for example, sham stimulation and concurrent cortical fMRI) to better isolate TMS-evoked spinal responses and more robustly delineate the causal pathways engaged by stimulation.

In this study, we introduced a multimodal framework that combined cortical TMS with spinal cord fMRI to map spinal circuit responses to non-invasive brain stimulation in vivo. In healthy participants, we found graded, intensity-dependent spinal BOLD responses that parallel motor output, demonstrating that the spinal cord exhibits dynamic and organized activity in response to cortical stimulation. By complementing electrophysiological measures of spinal excitability with spinal fMRI, this approach enables segment- and pathway-resolved imaging of descending motor control, highlighting activation within expected territories such as the left lateral corticospinal tract and ventral horn at C7–T1, during high-output sessions. To our knowledge, this is the first demonstration that concurrent TMS–SCfMRI can visualize how cortical drive is expressed within specific spinal segments and tracts in humans with high spatial specificity, establishing a foundation for probing sensorimotor circuits at the spinal level. Extending this framework to clinical populations could help identify residual functional pathways and guide individualized neuromodulation and rehabilitation strategies targeting the brain–spine axis.

## Acknowledgements

We thank Loan Mattera, Roberto Martuzzi, and Oliver Reynaud for their help with the acquisitions and setup. This study was partly funded by the Defitech Foundation (Morges, CH) to F.C.H, the Wyss Center for Bio and Neuroengineering (Lighthouse Partnership for AI-guided Neuromodulation) to F.C.H., and the Swiss National Science Foundation (205321− 207493) to D.V.D.V. This study was also supported by the MRI Platform of the Fondation Campus Biotech Geneva, Geneva, Switzerland co-founded and supported by EPFL, University of Geneva, and Geneva University Hospitals. Certain parts in the figures were created using biorender.

## Author Contributions

E.R., F.C.H., and D.V.D.V. initiated the study. E.R., R.J., and F.C.H. designed the protocol and collected the dataset. E.S. processed and analyzed the data. E.S. and D.V.D.V. wrote the paper.

## Data and Code Availability

Data and codes will be made available at the OSF repository.

## Supplemental Information (SI) for

### Simulated E-field spatial distributions

Figure. S1 depicts the simulated electrical field (E-field) spatial distributions across three TMS intensities (80% MT, 100% MT, and 120% MT) for a representative subject.

### TMS-evoked potentials at 120% (MT)

Figure. S2 depicts the TMS-evoked potentials at 120% motor threshold (MT) for a representative subject from High-output group and Low-output group in session-2.

### Data Quality

Figure. S3 depicts data quality assessment.

### TMS-intensity Dependent Effects on Spinal Cord BOLD Signal

To investigate whether spinal cord BOLD responses varied systematically with different TMS intensities, we analyzed activation maps derived from the statistical analysis detailed in Section 2.7-A2 in HOG of Session-1. Group-level results revealed a clear, intensity dependent increase in the proportion of active spinal voxels (*p* < 0.005, FWE-corrected) (Figure S4 A). The average proportion of active voxels was 0.0 ± 0.0 for the Low condition, 9.79 ± 6.42 for Medium, and 49.98 ± 17.74 for High intensity. In terms of spatial distribution, no significant activation was observed in the Low condition. For the Medium condition, activation extended into the contralateral ventral horns and intermediate zone at C6–T1, with additional dorsal activity at C6 and C8. The High condition exhibited the most widespread activation, involving the dorsal columns across C7-T1 spinal levels, with bilateral dorsal horns activation at C7 level, ipsilateral dorso-ventral activation at C8-indicating robust bilateral engagement of both sensory and motor pathways. At C6 level activation was localized to contralateral dorsal horn and ipsilateral motor horn. At C5 level, the activation was largely localized to dorsal columns and ipsilateral hemicord. In addition, parameter estimates (extracted as defined in 2.7) did not show any significant increase (*p* < 0.05) from Low-Medium, Medium-High, and Low-High conditions, across all spinal levels, except C6, where a significant increase was observed between Medium-High condition (*p* = 0.026). This observation suggests that, contrary to Session-2, the gray matter activations in Session-1, failed to show a significant intensity-dependent activation increase across spinal levels.

### Effect of Modulation on Spinal Cord BOLD Signal

To assess whether trial-by-trial variations in MEP amplitudes were associated with corresponding fluctuations in spinal BOLD responses, we analyzed activation maps derived from the MEP-based parametric GLM described in Section 2.7-A3 in HOG of Session-1. Group-level statistical maps were generated for both regressors using non-parametric permutation testing with TFCE approach. The unmodulated regressor showed activation (*p* < 0.05, FWE-uncorrected) spanning C5 to T1 spinal levels, with distributed spatial extent across white-matter and gray-matter regions (Figure S5). The modulated regressor, however, showed activation (*p* < 0.05, FWE-uncorrected) primarily localized to the gray-matter regions at C5– C6 spinal levels while at C8–T1 spinal levels, it was largely localized to white-matter regions, particularly descending tracts. This holds consistent with the hypothesis of TMS-driven modulation in descending motor pathways.

---

**Figure S1:**
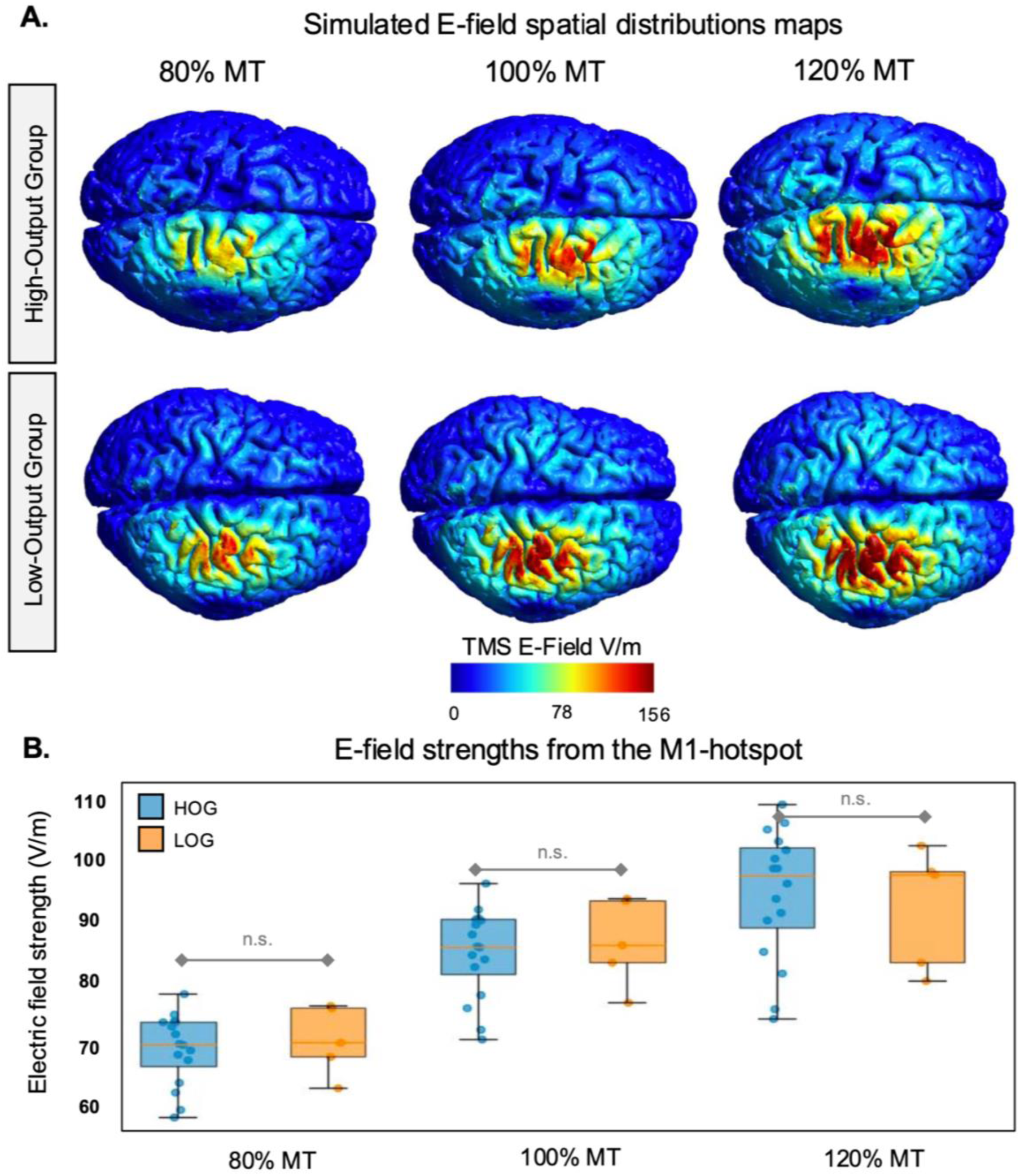
(A) Simulated electric field (E-field) spatial distributions generated by TMS at 80% MT (left), 100% MT (middle), and 120% MT (right) in a representative subject from HOG (top) and LOG (bottom). (B) Comparison of E-field strength in the M1-hand area, the intended stimulation target, at 80% MT (left), 100% MT (middle), and 120% MT (right). n.s.: *p* > 0.05.

**Figure S2.**
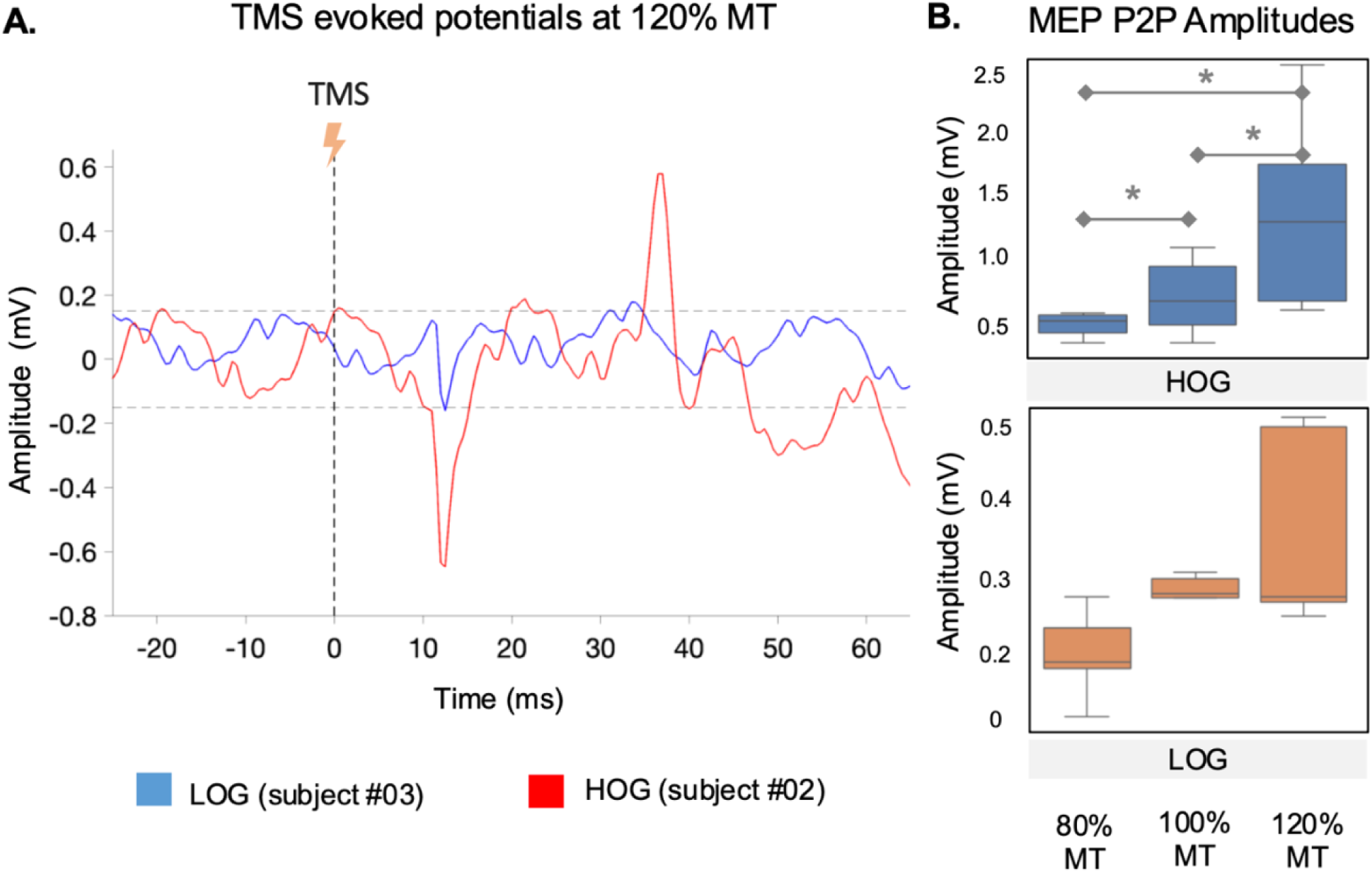
Example EMG responses and group-level MEP amplitudes in High Output and Low Output Group. (A) TMS-evoked EMG traces from the left FDI muscle at 120% MT for a representative HOG subject (top) and LOG subject (bottom) in Session-2, shown from −20 to 60 ms relative to the TMS pulse (0 ms). The setup-specific baseline EMG noise band (± 0.15 mV), estimated 100 ms pre-stimulus, is indicated by horizontal dashed lines. In the HOG example, a clear MEP deflection is visible between 30-40 ms and exceeds the noise band; in the LOG example, no discernible MEP is observed within this window. (B) Peak-to-peak MEP amplitudes across Low (80% MT), Medium (100% MT), and High (120% MT) intensities for HOG and LOG in Session-2, illustrating that MEPs are present in both subgroups but are consistently larger (>0.5 mV) and increase with intensity only in HOG, whereas LOG amplitudes remain close to baseline (0–0.5 mV). Asterisks denote significant differences (*: *p* < 0.05)

**Figure S3:**
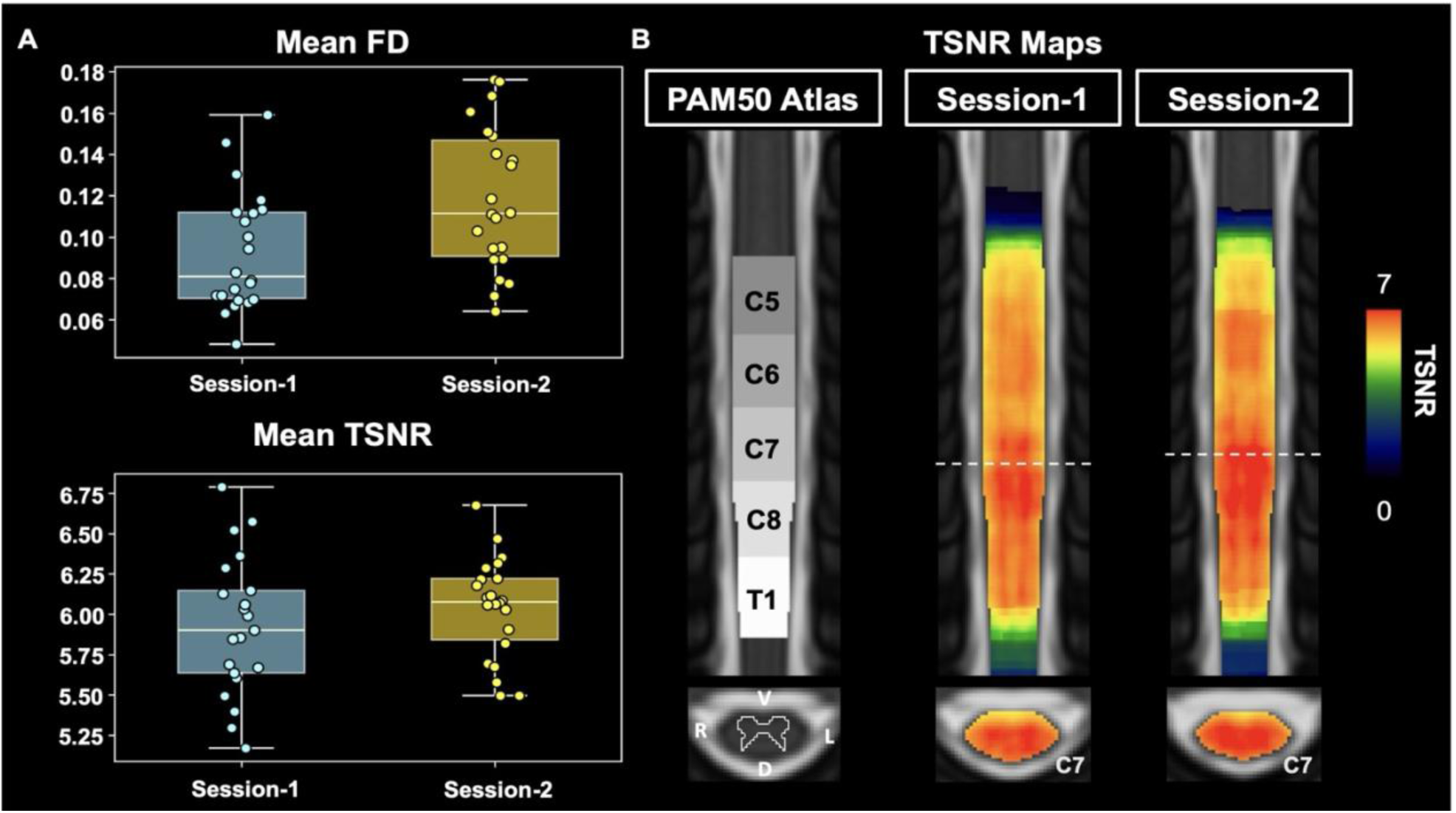
Data quality assessment.(A) Framewise displacement, and (B) TSNR maps.

**Figure S4:**
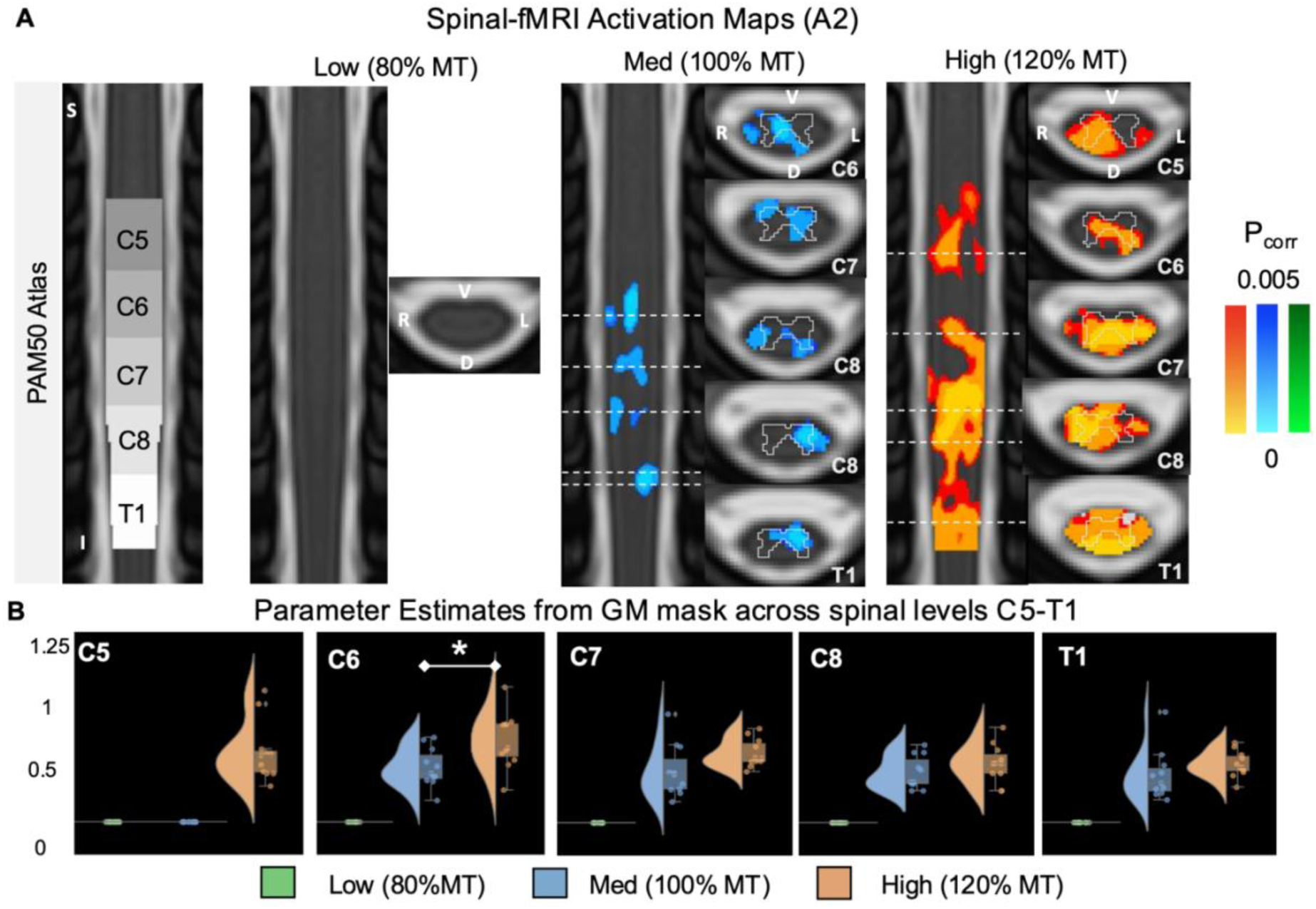
Intensity-Dependent Spinal BOLD Responses to TMS. (A) Group-level spinal activation maps for three TMS intensities (Low, Medium, High), thresholded at *p* < 0.005 (FWE-corrected). Axial slices are overlaid on the PAM50 template and labeled by spinal level (C5–T1). The dotted lines on coronal views marks the represented axial location. S = superior, I = inferior, L = left, R = right, D = dorsal, V = ventral. (B) Parameter estimates from activated voxels within the gray matter across spinal levels (C5–T1) for Low, Medium, and High TMS intensities, illustrating intensity-specific BOLD response amplitudes across spinal levels.

**Figure S5:**
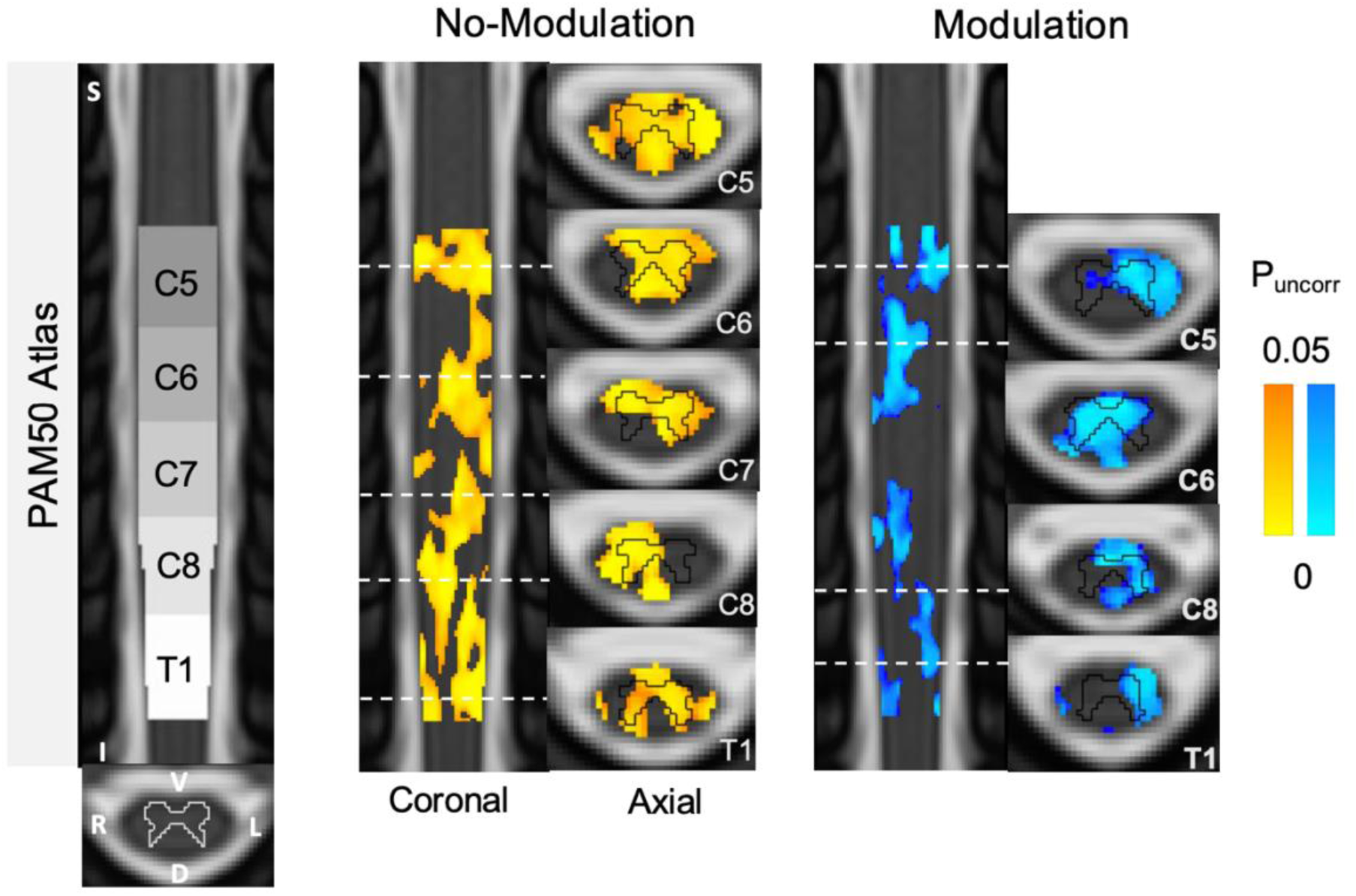
Trial-by-Trial Modulation of Spinal BOLD Signal by MEP Amplitudes. Group-level activation map for the unmodulated regressor (reflecting trial-wise TMS events) shows distributed activation spanning C5–T1, across both gray and white matter regions, thresholded at *p* < 0.05, FWE-uncorrected. The dotted lines on coronal views marks the represented axial location. S = superior, I = inferior, L = left, R = right, D = dorsal, V = ventral.

